# The idiosyncratic nature of confidence

**DOI:** 10.1101/102269

**Authors:** Joaquin Navajas, Chandni Hindocha, Hebah Foda, Mehdi Keramati, Peter E Latham, Bahador Bahrami

## Abstract

Confidence is the ‘feeling of knowing’ that accompanies decision making and guides processes such as learning, error detection, and inter-personal communication. Bayesian theory proposes that confidence is a function of the probability that a decision is correct given the evidence. Empirical research has shown, however, that humans tend to report confidence in very different ways. This idiosyncratic behaviour suggests that different individuals may perform different computations to estimate confidence from uncertain evidence. We tested this hypothesis by collecting confidence reports from healthy adults making decisions under either visual or numerical uncertainty. We found that for most individuals, confidence did indeed reflect the perceived probability of being correct. However, in approximately half of them, confidence also reflected a different probabilistic quantity: the observed Fisher information. We isolated the influence of each of these two quantities on confidence, and found that this decomposition is stable across weeks, and consistent across tasks involving uncertainty in both perceptual and cognitive domains. Our findings provide, for the first time, a mechanistic interpretation of individual differences in the human sense of confidence.

## Introduction

Understanding the computational basis of individual differences in human cognition has fundamental implications for medical and biological sciences as well as for economics and social sciences. A prime example is confidence, which plays a key role in a wide range of aspects in life, including learning to make better decisions^1^, monitoring our actions^2^, cooperating effectively with others^3, 4^, and displaying good political judgment^5^. One of the most intriguing features of confidence is that humans tend to communicate this feeling in a largely idiosyncratic way: although confidence reports are typically stable within each person, they tend to be variable across the population^6, 7^. For instance, different individuals performing the same task generate distributions of confidence ratings with different mean and shape^7^. In addition, the correlation between confidence and objective performance varies for different people, and is related to individual variations in brain structure^8^ and connectivity^9, 10^. While a vast literature has focused on the biological *correlates* of individual differences in human confidence^8––10^, the computational *roots* of this phenomenon remain unclear.

Previous research in sensory psychophysics^8, 11^ and value-based decision making^10^, assumed that confidence is a function solely of the perceived probability of being correct. This assumption is reasonable: confidence should only reflect this subjective probability^12–14^. However, based on this normative framework, previous studies explained differences between people as measurement noise^15^, or as individual differences in the ability to reflect the probability of being correct^8, 9^. This may have been an oversimplification: there is extensive literature showing that confidence can be influenced by factors other than the probability of being correct^16^, such as the reliability of sensory stimuli^2, 13^, the magnitude of sensory data^11^, post-decisional biases^17^, and even personality traits^7^.

Here we set out to determine what probabilistic quantities, besides perceived probability of being correct, contribute to individual differences in human confidence. We focused on a categorical task, in which subjects had to decide whether the average of a set of items was above or below a decision boundary, and then report their confidence. For about half of the subjects, confidence did depend solely on the perceived probability that they were correct. However, for the other half, confidence also depended on a different statistical quantity: the inverse variance, which we refer to as the observed Fisher information^18, 19^ (to avoid ambiguity associated with other terms used previously such as “certainty”^14^ or “precision”^1^). Moreover, the dependence of confidence on the perceived probability of being correct and the Fisher information was stable across experiments performed weeks apart. Finally, we show that the dependence of confidence on the perceived probability of being correct was stable across tasks involving uncertainty in the perceptual and cognitive domain, but the dependence on Fisher information was not. This is consistent with the predictions of a recent theoretical account arguing that Fisher information is encoded by domain-specific neural populations^14^. Overall, these findings provide a mechanistic interpretation of individual differences in the human sense of confidence.

## Results

In the perceptual task (Experiment 1), participants observed a sequence of 30 tilted Gabor patches presented at the fovea in rapid (4 Hz) serial visual presentation (**Fig.1a**). At the end of the sequence, participants decided whether the mean orientation of the patches was clockwise or counter-clockwise relative to vertical. Participants then reported how confident they were in their decision on a scale from 1 to 6. To manipulate uncertainty, we pseudo-randomly drew the orientation samples from uniform distributions with exactly the same mean (+3 degrees or -3 degrees) but different variances on different trials (**Fig. 1b**). Participants performed better as variance decreased (**Fig. 1c**, one-way repeated measures ANOVA, F_(3,29)_=231.4, *p*<10^-10^).

To fit the choices of each participant, we assumed that they keep track of the mean orientation, which they update after each stimulus presentation. To update their estimate of the mean, we considered a model in which participants combine a noisy estimate of the current sample with their previous estimate of the mean,

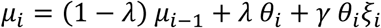
where *μ_i_* is the estimate of the mean after *i* samples (*μ_0_* = 0), 0 < λ < 1 determines the relative weighting of recent versus more distant samples, *θ_i_* is the actual orientation of the *i*^*th*^ sample in the sequence, *ξ_i_* is sampled from the standard normal distribution, and *γ* is a free parameter indicating the strength of the noise. The multiplicative nature of the noise ensures that the uncertainty in the update of the estimate scales with the size of the observed sample, *θ_i_*, as has been observed in numerous domains, including visual perception^20^, numerical cognition^21^, and the perception of time^22^. At the end of the sequence, choice is determined by the sign of the final value of the mean (*μ*_30_): the agent chooses clockwise if *μ*_30_ is positive, and counter-clockwise if *μ*_30_ is negative.

We also tested an alternative model that tracks the mean of the sequence in a deterministic way, and then makes stochastic decisions. This model, however, failed to explain the trend in **Fig. 1c**, which shows that performance increases as variance decreases (see **Supplementary Fig. 1** for details and model comparison).

**Figure 1.**
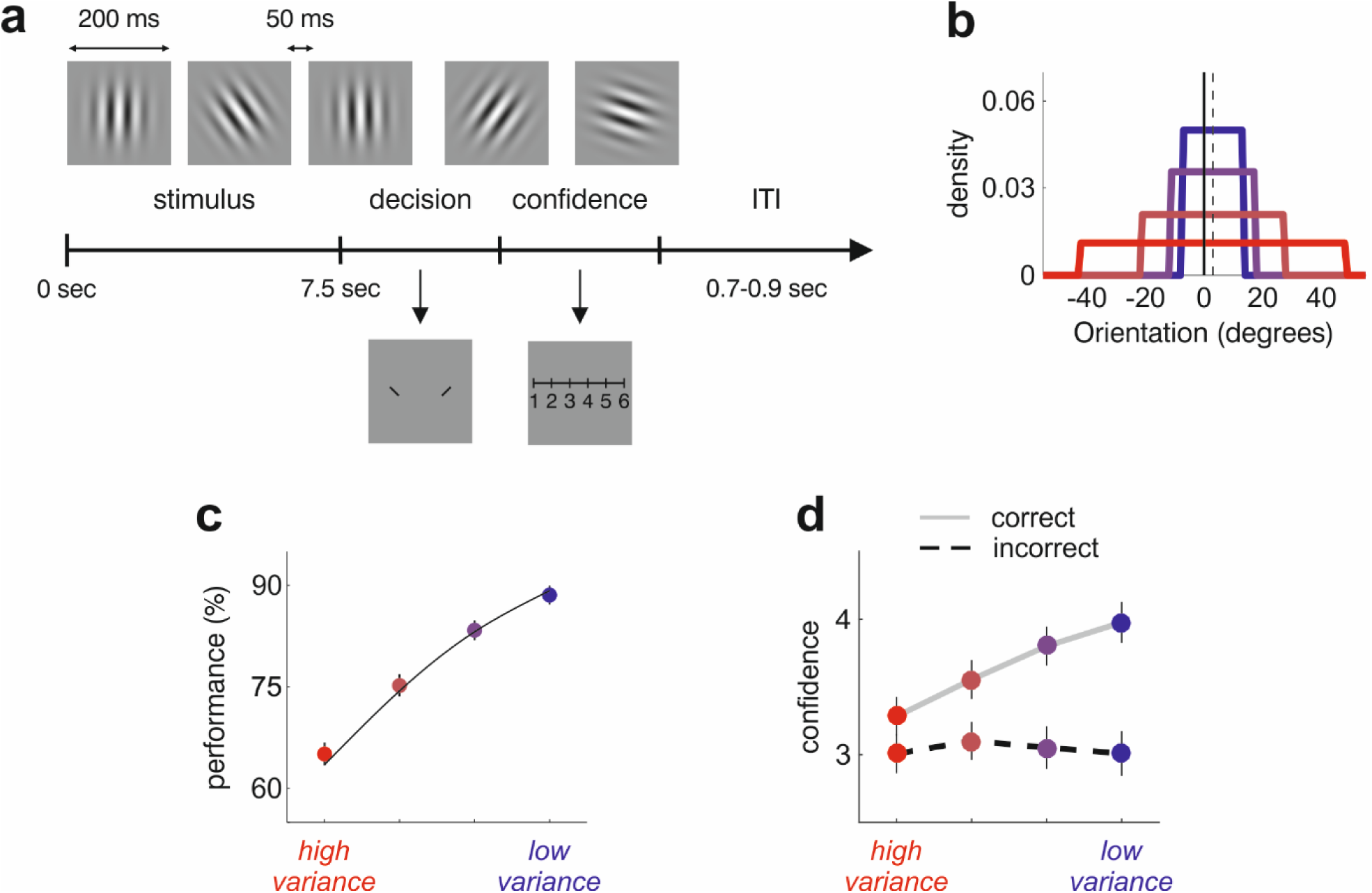
Tracking mean evidence in rapid serial visual presentations. **(a)** 30 tilted Gabor patches were serially flashed at the fovea, updated at 4 Hz. Participants made a binary decision about whether the mean in the sequence was tilted to the right or left, followed by a confidence rating. Full details of the task are available in Online Methods. **(b)** The samples were drawn from a uniform distribution with mean, *m*, set to either exactly +3 degrees or exactly −3 degrees. The dashed line shows *m*=+3. The endpoints of the uniform distributions were *m*±*v*, with *v* = 10, 14, 24, or 45 degrees, yielding four conditions with four different variances. **(c)** Performance increased with decreasing variance. Dots show the grand-average performance, and vertical lines depict the s.e.m. The solid black curve shows the best fit of the stochastic updating model (Equations [1] and [2]). **(d)** Confidence reports averaged over all subjects. Vertical lines show s.e.m. At the population level, confidence in incorrect trials remains approximately constant as a function of variance.

### Computation of confidence

To compute confidence, participants need to have an estimate of the variance of *μ*_30_. We assumed that they are able to compute the true variance associated with Equation [1]. Thus, perceived variance, denoted 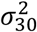, is given by

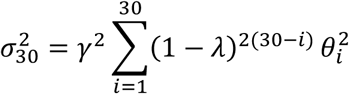
The model described by Equations [1] and [2], which we call the stochastic updating model, is illustrated in the left two panels of **Fig. 2a**. Given *μ*_30_ and 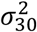, subjects can compute, on each trial, the perceived probability of being correct, 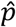(correct) (shaded area under the Gaussian distribution in **Fig. 2a**). Participants can also compute the Fisher information using 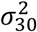 (top right panel in **Fig. 2a**).

**Figure 2.**
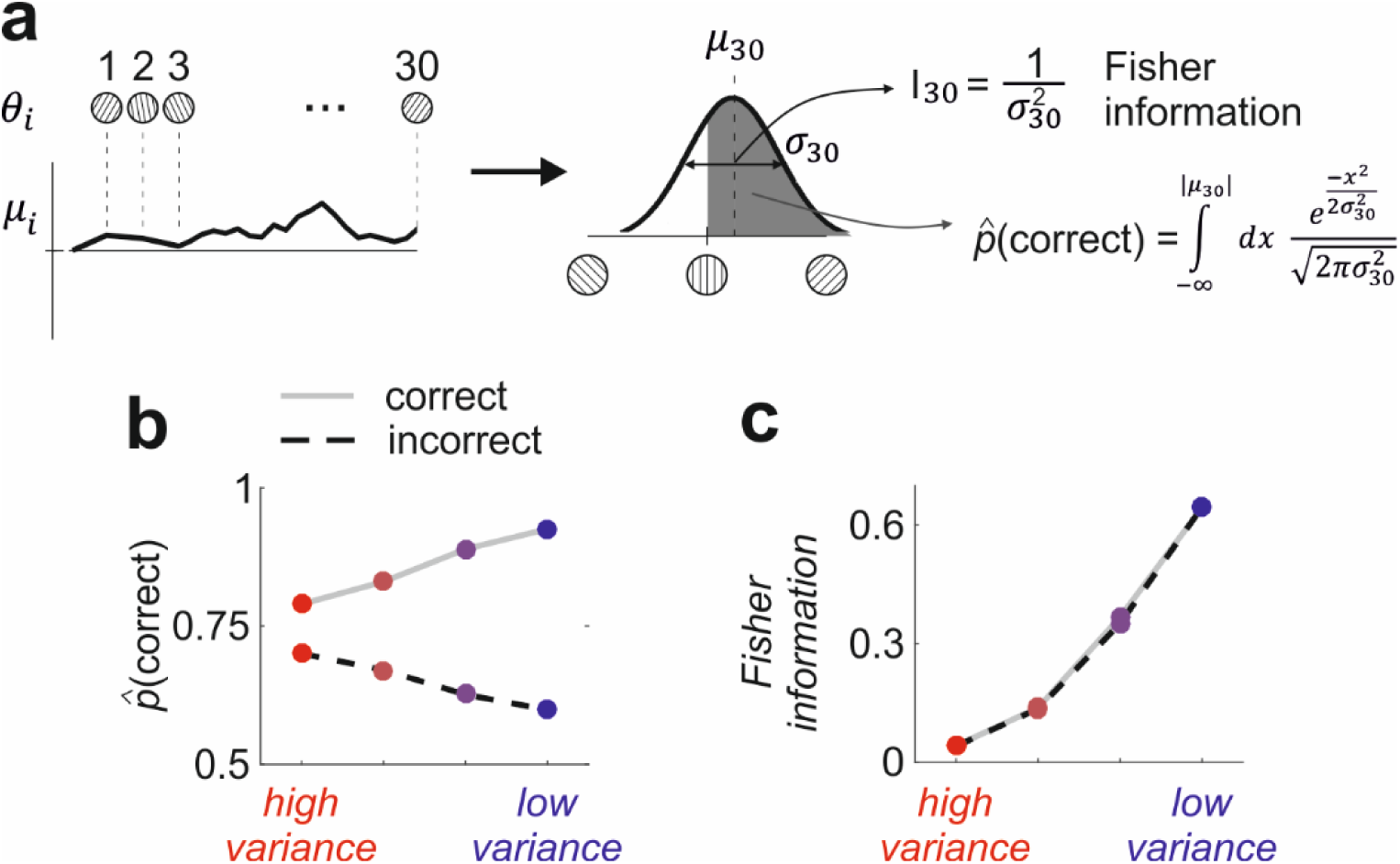
Estimating confidence. (**a**) Each trial consists of 30 presentations of tilted Gabor patches. At each presentation(*θ*_*i*_) the mean (*μ*_*i*_) is updated by combining the estimate on the previous sample with a noisy version of the current Gabor patch. The black line represents one realisation of the model. At the end of the sequence, the subject makes a decision based on the sign of *μ*_30_. The subjective probability of being correct and the observed Fisher information are then computed according to the equations shown in the right panel; see Online Methods for full details. (**b**) The perceived probability of being correct, 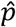(correct), averaged over variance condition for correct trials (solid grey line) and incorrect trials (dashed black line), and also averaged across participants. For correct trials, this quantity increases with decreasing variance (solid grey line); for incorrect trials it shows the opposite pattern (dashed black line). (**c**) Observed Fisher information increases both for correct and incorrect trials (same markers as panel **b**).

Using this model, we estimated the expected values of 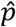 (correct) and Fisher information for different variance conditions, separated by correct and incorrect trials (see Online Methods, Equation [9], and **Fig. 2b,c**). On correct trials, both quantities increase with decreasing variance (solid grey lines in **Figs. 2b** and **2c**), as does confidence (**Fig. 1d**). If we had access only to correct trials, we would not know whether confidence was influenced by perceived probability of being correct or Fisher information. However, on error trials, these two quantities show opposite trends: 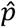(correct) decreases with decreasing variance while Fisher information increases (dashed black lines in **Figs. 2b** and **2c**). Confidence in errors, on the other hand, was relatively independent of variance (F_(3,29)_=0.57, *p*=0.63). One explanation of this phenomenon is that participants base confidence on both their perceived probability of being correct and, as a heuristic, the observed Fisher information. If confidence were an increasing function of both 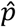(correct) and Fisher information, and subjects weighted them approximately equally, confidence could be a relatively flat function of variance on incorrect trials (**Fig. 1d**), and at the same time could increase with decreasing variance on correct trials.

The analysis presented so far is based on population-averaged data (**Fig. 1d**), so it is uninformative about differences among individuals. To determine whether, and how, 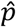(correct) and Fisher information influence confidence within subjects, we looked at the data of each individual. As expected^6, 7^, we observed substantial inter-individual differences (**Fig. 3**). Some subjects did indeed base confidence solely on 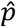(correct). However, in approximately half of them, confidence appeared to be influenced – at least to some degree – by Fisher information. To quantify this, we regressed^23^ confidence reports against model-based estimates of 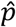(correct) and information. **Fig. 3
** shows a scatter plot of the regression weights for 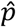(correct) and Fisher information. In 13 out of the 30 participants, confidence significantly reflected 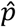(correct) but not information. In 14 other participants, however, confidence significantly reflected both 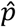(correct) and information. One participant’s confidence conveyed only information but not 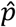(correct), and finally, for two participants, confidence did not reflect either of the two quantities.

**Figure 3.**
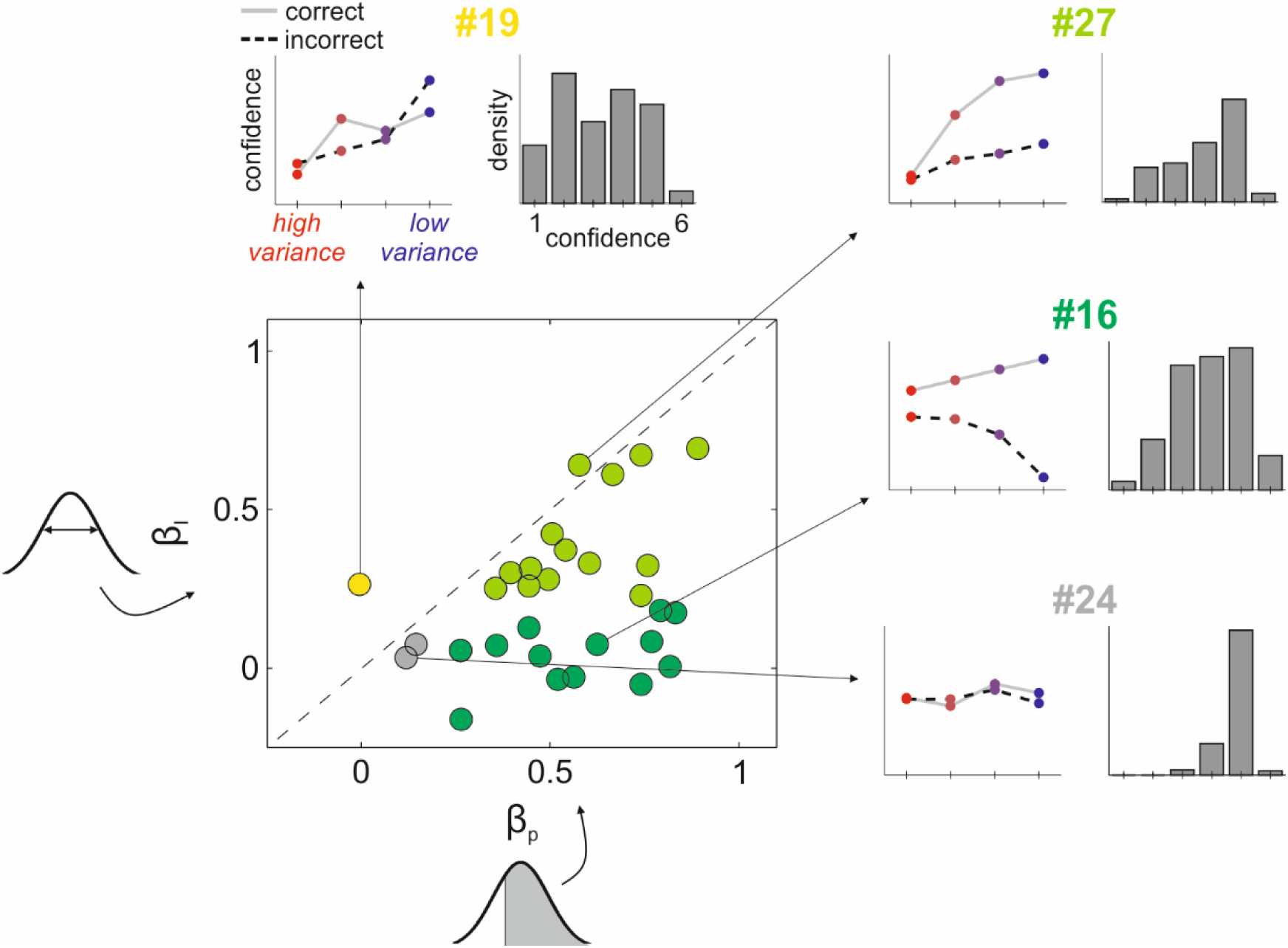
Analysis of confidence across individuals. The main panel in the lower left shows regression weights on confidence for different individuals. x-axis: weight of the probability of being correct (*β*_*p*_); y-axis: weight of information (*β*_*I*_). Each dot is a different participant, and the colour codes for significance (at the 0.05 level) as follows: dark green, only *β*_*p*_ was significant; light green, both *β*_*p*_ and *β*_*I*_ were significant; yellow, only *β*_*I*_ was significant; grey, neither was significant. Insets along the top and right margins show average confidence and confidence distributions for four representative participants. Left plots: mean confidence across different variance conditions, split by correct (solid grey line) and incorrect (dashed black line) trials. Right plots: probability distribution over confidence. For participant #19 (yellow dot), confidence reflected only information: confidence increased with variance for incorrect trials. For participant #16 (dark green dot), confidence reflected only the perceived probability of being correct: confidence in error trials decreased with increasing variance. For participant #27 (light green dot), confidence reflected a mixture of both computations. For participant #24 (grey dot), confidence was not modulated by either of these quantities.

### Stability across time

The regression model identified seven parameters for each individual (see Online Methods, Equation [10]): a weight for 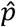(correct), denoted; *β*_*p*_ a weight for information, denoted; *β*_*I*_ and five parameters *α*_*j*_ (*j* = 1,…,5). The latter are the average log odds of observing a confidence rating greater than *j*; from these we selected the mid-value, *α* _3_, which is based on splitting the confidence scale in halves. The parameter *α*_3_ was correlated with the average confidence across the entire experiment (*r*=0.84, *p*<10^−8^), and so indicates how under- or overconfident a given participant is; we thus refer to *α*_3_ as the average confidence. We confirmed that individual differences in these parameters(*β*_*p*_, *β*_*I*_, and *α*_3_) are not simply explained by how well our model fit decisions (see Online Methods). The regression weights, *β*_*p*_ and, *β*_*I*_ were correlated with measures of confidence used in previous studies: *β*_*p*_ was correlated with how well confidence predicted accuracy, and *β*_*I*_ was correlated with the subjects’ ability to discriminate different variances (see Online Methods). The three selected variables were uncorrelated with each other across the population (*r*<0.35, *p*>0.1 for all pairwise comparisons between *β*_*p*_, *β*_*I*_, and *α*_3_).

This analysis would be no more than a model-fitting exercise if a different profile – that is, a different relationship between confidence, 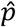 (correct), and Fisher information – emerged when the same participants were retested. To test for stability, in Experiment 2 we retested 14 of the participants from Experiment 1 approximately one month later. We observed that the three variables (*β*_*p*_, *β*_*I*_ *α*_3_) were correlated across experiments (**Fig. 4**), indicating that this decomposition is stable across time and informative of the identity of the participants. To further validate this observation, we found that the distance in the 3-dimensional space defined by(*β*_*p*_,*β*_*I*_ *α*_3_) within participants (across the two experiments) was smaller than the distance between different participants within an experiment (Wilcoxon rank sum test, *z*=4.0, *p*<10^−4^).

**Figure 4.**
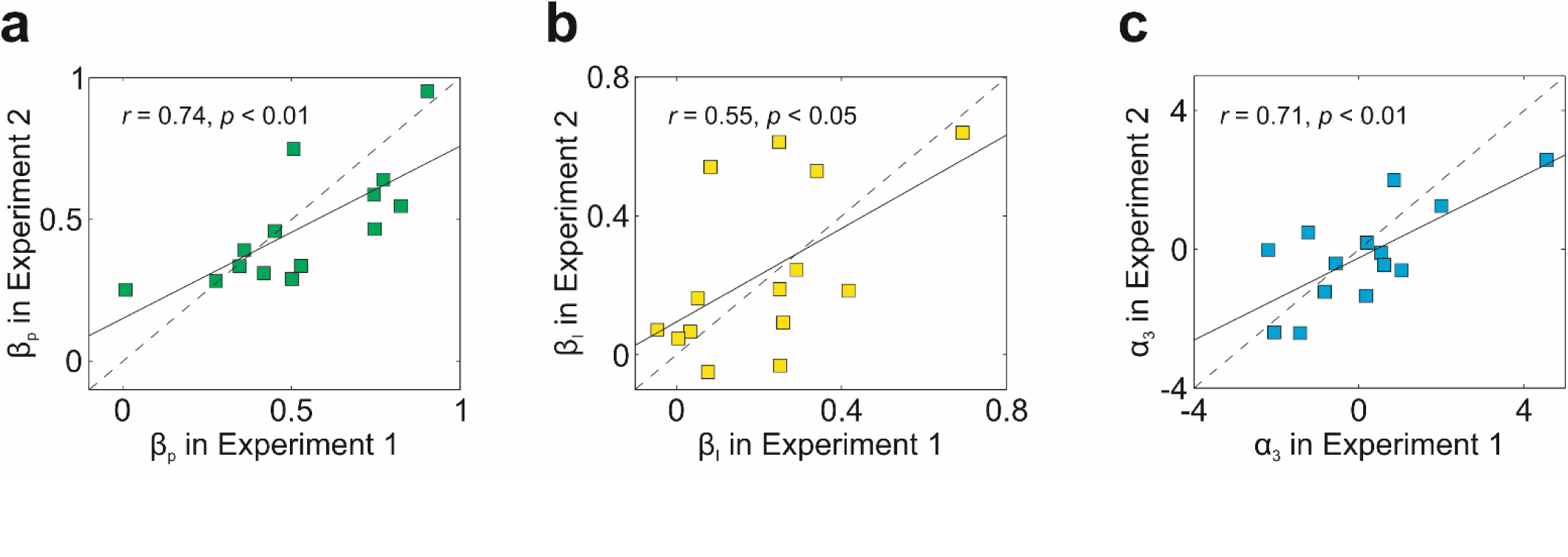
Stability across time. 14 participants of Experiment 1 were retested about a month later (35.2±2.4 days; range = 23-49 days). We probed stability by asking how much our three parameters (*β*_*p*_,*β*_*I*_ and *α*_3_) changed across experiments. (**a-c**) Correlation across experiments for *β*_*p*_ (**a**), *β*_*I*_ (**b**), and *α*_3_ (**c**). Each square is a different participant, the dotted line is the identity, and the value of *r* given in each box is the Pearson correlation coefficient. The three variables were significantly correlated across experiments, suggesting that this decomposition is stable across time. A non-parametric method to measure rank correlation across experiments yielded similar results (Spearman’s rank correlation, *r_s_*=0.82, *p*<0.001 for *β*_*p*_ *r_s_*=0.54, *p*<0.05 for *β*_*I*_, and *rs*=0.55, *p*<0.05 for *α*_3_).

### Consistency across tasks

To determine whether subjects compute confidence the same way across tasks – that is, whether they give the same weight to 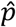(correct) and Fisher information, and have the same average confidence – we repeated our experiments on a cognitive task: averaging a sequence of numbers. In Experiment 3, a new group of 20 participants performed, in counterbalanced order, the visual task described above and a numerical averaging task (**Fig. 5**). In the numerical task, we presented two-digit numbers, updated at the same rate as in Experiment 1 (4 Hz). The task was to decide whether the mean of the sequence was greater or smaller than 50. Uncertainty was manipulated in the same way as in Experiment 1, but using different ranges to ensure comparable performance across tasks (see Online Methods for details).

In both tasks, accuracy increased with decreasing variance (**Fig. 5a,b**). A two-way repeated measures ANOVA with factors “variance” and “task” showed a significant main effect of variance (F_(3,19)_=194.3, *p*<10^−10^) but a non-significant effect of task (F_(1,19)_=2.5, *p*=0.13) or interaction (F_(3,19)_=0.84, p=0.47). Importantly, replicating Experiment 1, variance did not modulate confidence in error decisions (F_(3,19)_=0.2, *p*=0.89 for the visual task; F_(3,19)_=1.1, *p*=0.4 for the numerical task). Confidence in the visual task was not statistically different from confidence in the numerical task (F_(1,19)_=1.58, *p*=0.22, **Fig. 5c,d**). Decisions in both tasks were better fit by the stochastic updating model (Equations [1] and [2]) than the same alternative model we considered in the visual task (log-likelihood of the difference against zero: t_(19)_=5.2, *p*<10^−4^ for the cognitive task; t_(19)_=6.4, *p*<10^−5^ for the perceptual task), and our model-based analysis of confidence found similar results to Experiment 1 (see **Supplementary Figs. 2** and **3**).

**Figure 5.**
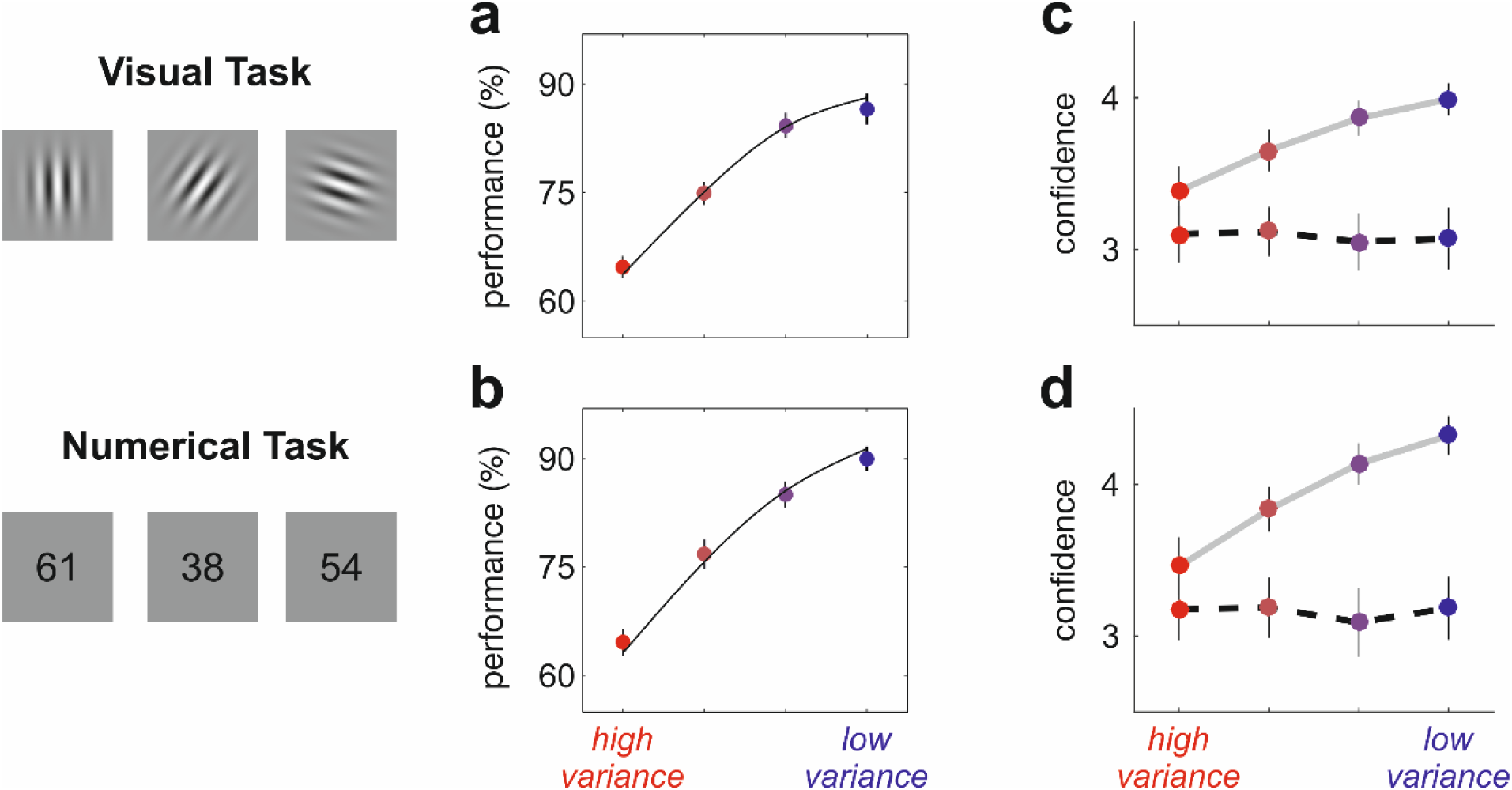
Decisions and confidence in Experiment 3 (N=20). (**a**,**c**): Visual task (replication of Experiment 1 with different participants; panel **a** corresponds to **Fig. 1c** and panel **c** to **Fig. 1d**). (**b**) Same as (**a**), but for the numerical task. (**d**) Same as (**c**), but for the numerical task. The similarity between panels **a** and **b**, and between panels **c** and **d**, indicate that, at least on average, the visual and numerical tasks lead to remarkably similar behaviour, despite the fact that one is perceptual and the other is cognitive.

We asked if our three regressors were consistent across the numerical and visual tasks. The within-participants distance in the 3-dimensional space was smaller than the between-participants distance (Wilcoxon rank sum test, *z*=3.3, *p*<0.001). The contribution of 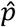(correct), *β*_*p*_ (*r*=0.74, *p*<0.001) and trait confidence *α*_3_ (*r*=0.63, *p*<0.01) were significantly correlated across tasks. However, the weights of information,*β*_*I*_ were uncorrelated across tasks (*r*=0.20, *p*=0.37), indicating that Fisher information has quantitatively different effects on confidence in visual and numerical tasks (**Fig. 6**). This result is in agreement with a recent theoretical account arguing that the inverse variance is represented by domain-specific neural populations^14^ (see Discussion).

**Figure 6.**
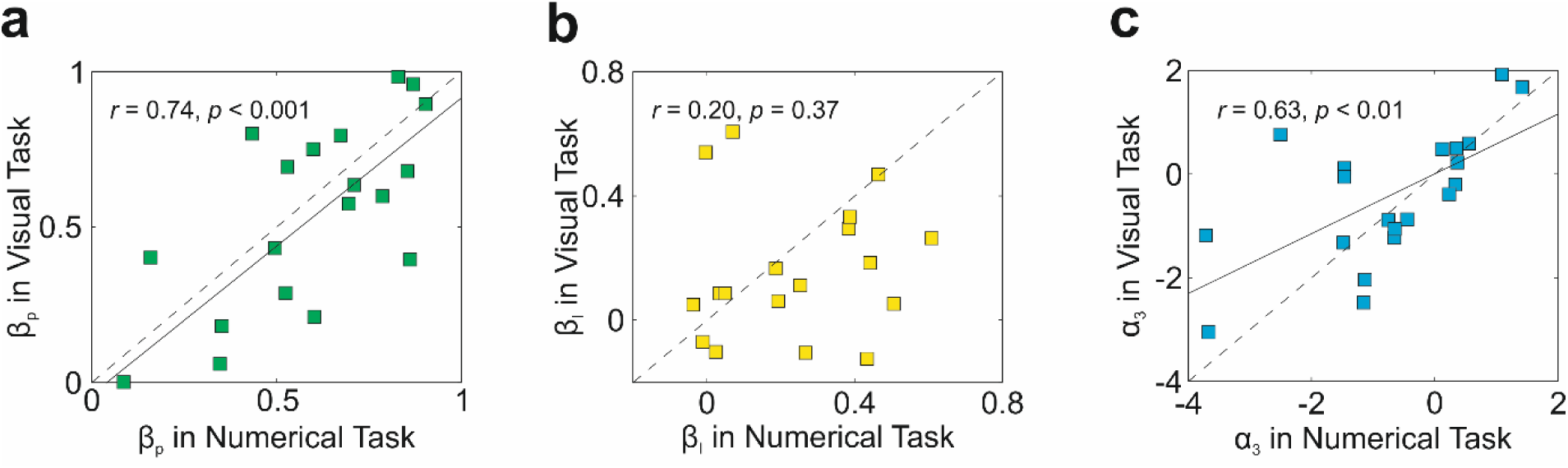
Consistency across tasks involving uncertainty in the perceptual and cognitive domain. 20 participants that were not tested in Experiments 1 or 2 performed one visual and one numerical task (Experiment 3). As in **Fig. 3**, we decomposed confidence in terms of the weight of 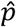(correct)(*β*_*p*_), the weight of information (*β*_*I*_), and the average confidence (*α*_3_). (**a-c**) Correlation across tasks for *β*_*p*_ (**a**), *β*_*I*_ (**b**), and *α*_3_ (**c**). Each square is a different participant, the dotted line is the identity, and the value of *r* given in each box indicates the Pearson correlation coefficient.*β*_*c*_ and *α*_3_ were positively correlated across tasks; however, the weights of Fisher information,*β*_*I*_ were uncorrelated across tasks. A non-parametric method to measure the correlation across experiments yielded similar results (*r_s_*=0.68, *p*<0.01 for,*β*_*p*_ *r_s_*=0.22, *p*=0.35 for,*β*_*I*_ and *r_s_*=0.62, *p*<0.01 for *α*_3_).

## Discussion

The computations underlying confidence have attracted considerable attention over the last several years, in part due to recent developments in model-based approaches^12–14^ combined with neurophysiological recordings in non-human animals ^24–26^ and neuroimaging in humans ^8–10, 27^. The standard approach consists of fitting a model to the entire population and treating inter-individual variability as noise^11, 15^. However, if such individual differences are robust over time, and consistent across tasks^7^, then treating them as noise limits our understanding of the computational processes underlying confidence. Here we found that inter-individual differences in confidence ratings are meaningful in terms of their underlying computations. In particular, we found that different individuals used different weightings for two probabilistic quantities: their perceived probability of being correct, and their certainty in their estimate of the task-relevant variable^14^, the latter quantified by the observed Fisher information^18, 19^. We isolated the contribution of each of these two quantities to confidence, and measured, for each individual: 1) the influence of the perceived probability of being correct on confidence (*β*_*p*_), 2) the influence of Fisher information on confidence (*β*_*I*_), and 3) the participants’ average confidence (*α*_3_). All three variables were stable across several weeks (**Fig. 4**), and two of them (*β*_*p*_ and *α*_3_) were stable across different tasks – one in the perceptual domain; the other in the cognitive domain (**Fig. 6**).

Previous research has shown reliable individual differences in the mean and shape of the distribution of confidence ratings^6, 7^, and in the extent to which confidence predicts behavioural accuracy^7, 8^. These properties are believed to be idiosyncratic and correlate with individual variations in personality trait^7^, brain structure^8^, and resting-state functional connectivity^9^. For example, individual differences in the correlation between confidence and accuracy were systematically linked to a frontal network including the anterior prefrontal cortex, ventro-medial prefrontal cortex, and rostro-lateral prefrontal cortex^8, 10, 28, 29^. These findings were based on decisions in a wide range of contexts, including visual^8^ and value-based^10^ choices. Although these studies provided interesting insights about the brain regions that correlate with individual differences in confidence, none of them explicitly asked what probabilistic quantities influence this variability. In parallel, theoretical accounts have formalised a definition of confidence as the perceived probability of being correct^12–15^, but without looking at individual differences. Here, we provide the first empirical evidence that the idiosyncratic nature of confidence is due to differences in the computation of confidence; more specifically, different individuals place different weighting on the perceived probability of being correct and the observed Fisher information.

In categorical tasks like ours, confidence should depend solely on the probability of being correct. So why does Fisher information affect confidence? We speculate that Fisher information, which is a useful measure of uncertainty in continuous quantities^1, 2, 13^, could serve as a mental shortcut that provides a proxy for the probability of being correct. This shortcut is reasonable, as Fisher information correlates with performance accuracy in our experiments (**Figs. 1c** and **2c**). Previous research in our group showed that interacting dyads do take shortcuts: they communicate a confidence signal that is close to, yet different from, the probability of being correct^3^. Similarly, we previously found that confidence can reflect the magnitude of sensory data^11^, a choice-independent quantity that also correlates with behavioural performance. Our finding that a heuristic, such as Fisher information, modulates confidence judgements about categorical decisions is in line with these studies.

### Predictions for neural data

Because the probability of being correct is a dimensionless quantity, and is universal across different sources of uncertainty, it could be encoded by a domain-general circuitry – for instance, by neurons in the prefrontal cortex^8, 10, 28, 29^. In contrast, Fisher information is a quantity with dimension, and so is presumably encoded by domain-specific populations^14^. For example, in the case of the visual task, certainty could be represented by neurons in primary visual cortex that are tuned to orientation^30^; and indeed, sensory uncertainty can be decoded from activity in the visual cortex^31^. In the same manner, numerical certainty could be represented by neurons in the parietal cortex tuned to different numerical quantities^32^, although this has not yet been tested.

An implication of our findings is that neurons representing confidence should receive input both from populations encoding the perceived probability of being correct and from populations encoding Fisher information. Consequently, because of differences in functional connectivity (which are likely to arise during learning and development) different individuals should have different weightings for the perceived probability of being correct and Fisher information on confidence. This is exactly what we saw in **Fig. 3**. In addition, because the perceived probability of being correct is encoded by domain-independent populations and Fisher information by domain-specific ones, the influence of perceived probability of being correct on confidence should be stable across domains, while the influence of Fisher information should be domain specific. This is exactly what we saw n in **Figs. 6a** and **6b**, respectively. Based on these observations, we can make a strong prediction, one that could be tested with neuroimaging studies: the larger the influence of Fisher information on confidence, the stronger the correlation between activity in domain-specific areas and confidence.

## Conclusion

The value of investigating individual differences in human behaviour and cognition was first recognised in the psychological sciences, with a special interest in high-level aspects such as intelligence^33^ and personality^34^. More recently, technical advances in magnetic resonance imaging have made it possible to develop a cognitive neuroscience of individual differences^35, 36^. Findings include neural correlates of individual differences in motor behaviour^37^, visual perception^38^, mood^39^, social network size^40^, and confidence^8–10^. While these studies provide valuable insights into the neural basis of inter-individual differences in human cognition, the mechanisms responsible for such differences remain unknown. To overcome this limitation, the next challenge is to build a computational neuroscience of individual differences. A first step in this direction is to understand the computations performed by healthy adults leading to inter-individual variability in behaviour. Our study represents the first effort to model differences in human confidence, paving the way towards finding how these computations change under development^41^, aging^42^, and psychiatric disorders^43^.

## Online Methods

### Participants

50 healthy adults (aged 18-45, 43 right-handed, 31 female) with normal or corrected-to-normal vision participated in this study. All participants were recruited through advertisement at University College London, and gave written informed consent. We collected data from 84 experimental sessions lasting approximately 90 minutes each. Participants were paid £10 per hour. All experimental procedures were approved by the research ethics committee at University College London.

### Display

Stimuli were generated using the Cogent Toolbox (http://www.vislab.ucl.ac.uk/cogent.php) for MATLAB (Mathworks Inc). Participants observed an LCD display (21-inches monitor; refresh rate: 60 Hz; resolution: 1024 × 768 pixels) at a viewing distance of approximately 60 cm.

### Experiment 1: Visual task

30 participants performed Experiment 1, which consisted of an orientation averaging task (**Fig. 1**). Observers viewed a sequence of 30 tilted Gabor patches over a middle grey background (standard deviation of the Gaussian envelope: 0.63 deg; spatial frequency: 1.57 cycles deg^−1^; contrast: 25%) flashed in rapid succession at the centre of the screen. Each patch was presented for 200 ms with an inter-stimulus interval of 50 ms, resulting in an update rate of 4 Hz. Once the sequence finished, the participant was asked to judge whether the mean orientation of the patches was tilted clockwise or counter-clockwise relative to the vertical. The response alternatives consisted of two tilted lines presented in the left and right visual field (size: 2.2 deg, location: 11.3 deg left or right to the centre of the screen). The position of the response alternatives was randomly assigned and counter-balanced across trials. To select the option displayed in the left, participants pressed the ‘Q’ button of a QWERTY keyboard using the left hand; to select the option on the right, they pressed the ‘P’ button. Participants were then asked to report their confidence on a rating scale from 1 to 6. A horizontal line was presented at the centre of the screen (length: 18.9 deg) with 6 equally-spaced marks signalling different levels of confidence. Participants moved a cursor to the left or right of the scale by pressing the ‘Q’ or ‘P’ buttons respectively. The initial point in the scale was randomly chosen on a trial-by-trial basis. Once the participants selected a confidence rating, they pressed the space bar to continue. After an inter-trial interval (which was uniformly distributed between 0.7 and 0.9 seconds), a new trial began.

The orientations of the patches were drawn from uniform distributions with mean *m* and endpoints *m*±*v*. We used distributions with two different means (*m* = +3 or −3 degrees) and four different variances (given by their different endpoints: *v* = 10, 14, 24, or 45 degrees). Uniform distributions were pseudo-randomly sampled such that the mean was exactly ±3 degrees on every trial. Orientations were randomly shuffled to define the presentation order. The experiment consisted of 400 trials: 50 trials for each of the eight distributions. Blocked feedback was given every 20 trials by a message displaying the number of correct trials in that block. Each block comprised 5 trials of each variance condition presented in random order. Therefore, performance for different variance conditions could not be learned from feedback.

### Experiment 2: Stability across time

All participants of Experiment 1 were invited to perform the visual task a second time, approximately one month later. 14 participants accepted the invitation and were re-tested. Experiment 2 was performed 35.2±2.4 days after Experiment 1 (range: 23-49 days). Experimenters were blind to the results of Experiment 1 when testing participants in Experiment 2.

### Experiment 3: Stability across the perceptual and cognitive domain

20 healthy adults who did not participate in Experiment 1 or 2 performed Experiment 3. Participants performed two sessions: the visual task described in Experiment 1 and a numerical averaging task. Half of the participants performed the visual task first. The second session was performed 9.7±2.9 days (range: 1-27 days) after the first one. Experimenters were blind to the results of the first session when testing the participants in the second session.

The numerical task was identical in structure to the visual task but, instead of Gabor patches, two-digit numbers (size: 3.8 deg; font: Arial) were presented. The colour of the numbers (black or white over a middle grey background) was randomly chosen at each presentation. Participants were instructed to decide whether the mean of the sequence was greater or smaller than 50. Numbers were sampled from uniform distributions with mean *m* = 47 or *m* = 53, and endpoints *m*±*v* were defined by *v* = 8, 11, 16 or 22. These values were chosen, through pilot experiments with a different set of participants, to obtain performances similar to that observed in Experiment 1. Uniform distributions were pseudo-randomly sampled such that the mean of the sequence was exactly *m* on each trial. Decisions were collected in the same way as in Experiment 1: a response screen with two options (“smaller” and “greater”) was presented on both sides of the visual field. Participants gave their answer, and indicated confidence, using the same keys as in the visual task.

### Model fitting

To fit the stochastic updating model (Equations [1] and [2]) to the participants’ decisions, we find, for each individual, the parameters *λ* and *γ* that maximise the log likelihood, 
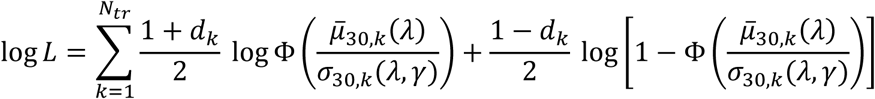
where Φ is the standard cumulative normal function, *d_k_* is the decision on trial *k*(+1 if clockwise, −1 if counter-clockwise), σ_30, *k*_(*λ, γ*) is obtained from Equation [2], and

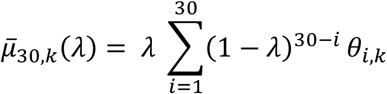
is the mean value of *μ*_30_ on trial *k*. (A minor technical point: Equation [4] describes the visual task; the cognitive task is the same except that the mean is offset by 50.)

### Estimating the Fisher information and the perceived probability of being correct

Based on the best fitting parameters *λ* and *γ* derived from the stochastic updating model, we estimated, on a trial-by-trial basis, the observed Fisher information and the expected perceived probability of being correct. The observed Fisher information is just the inverse variance of the participants’ estimate, the latter computed via Equation [2] (**Fig. 2a**). The expected perceived probability of having made a correct decision, *d*, is given by

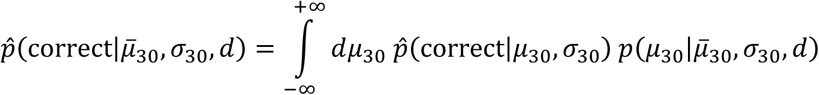
The first term inside the integral, 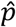(correct| *μ*_30_, *σ* _30_), is the shaded area under the Gaussian in **Fig. 2a**; consequently, it is given by the cumulative normal distribution,

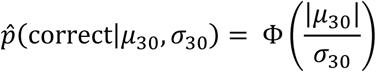

The second term in the integral, *p*(*μ*_30_|*μ̅*_30_, *σ*_30_, *d*), is the probability of observing *μ*_30_ given *μ̅*_30_, *σ*_30_, and, importantly, the decision, *d*. If the decision is clockwise (*d*=+1), *μ*_30_ must be positive, whereas if the decision is counterclockwise (*d*= −1), *μ*_30_ must be negative. We can take these constraints into account using the Heaviside step function, Θ(*x*) (which is 1 if *x*> 0 and 0 otherwise), yielding

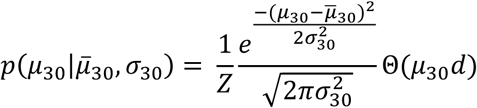
where *Z* is the normalisation constant,

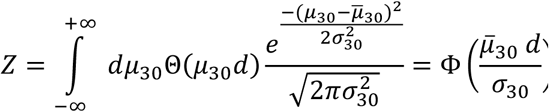

Combining these two expressions, we have

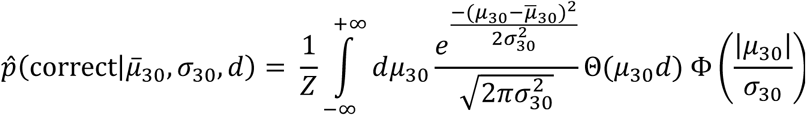

On each trial, 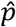(correct|*μ̅*_30_, *σ*_30_, *d*) was computed numerically using Matlab. Note that the expected perceived probability of being correct (Equation [9]) is dependent on the decision, *d*, whereas the Fisher information (Equation [2], **Fig.2a**) is choice-independent.

### Ordinal regression of confidence reports

We ran for each individual a multivariate ordinal regression^23^. For each of the five possible splits in the rating scale, this regression fits a logistic model with fixed effects and different offsets,

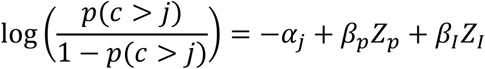
where 1 ≤ *j* ≤ 5, *c* denotes confidence, and *Z_p_* and *Z_I_* are z-scored estimates of the perceived probability of being correct and Fisher information on each trial. The outputs of this regression are the offsets *α*_1_,…, *α* _5_, and the weights *β*_*p*_ and *β*_*I*_ To summarise the computations underlying confidence, we selected *α*_3_ (the offset when splitting the scale in halves, which we refer to as the average confidence), *β_p_* (the weight of the probability of being correct on confidence) and *β_I_* (the weight of information on confidence).

### Does model fitting explain our findings?

We asked if individual differences in how well our model fitted decisions could explain inter-individual variability in the parameters, *β*_*p*_, *β*_*I*_ and *α*_3_. To do this, we correlated these values with the observed maximum log-likelihood,

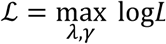
where log *L* is obtained using Equation [3]. We observed that ℒ was not significantly correlated with *β*_*I*_ (*r*=0.27, *p*=0.15) or *α*_3_ (*r*=0.26, *p*=0.15); however, we did observe a positive correlation between ℒ and *β*_*p*_(*r*=0.59, *p*<0.001). This could potentially mean that, for subjects with a low,*β*_*p*_ decisions were not well explained by our model (Equation [1]). Alternatively, because ℒ is positively correlated with the mean accuracy across the entire session (*r*=0.71, *p*<10^−4^), it could reflect that these participants behaved more randomly. To test this possibility, we computed the expected log-likelihood under the assumption of a perfect fit,

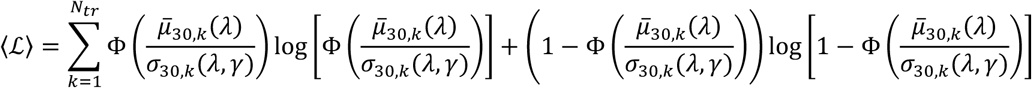
and measured the quality of the fit using the deviance, *D*= −2(2112;−〈ℒ〉). All three parameters were uncorrelated with *D* (*r*=0.22, *p*=0.24 for;*β*_*p*_ *r*=-0.12, *p*=0.54 for;*β*_*I*_ *r*=0.24, *p*=0.19 for *α*_3_), and *D* was uncorrelated with average performance (*r*=0.22, *p*=0.23). This indicates that individual differences in,*β*_*p*_*β*_*I*_ and *α*_3_ are not explained by inter-individual variability in the goodness of the fit.

### Comparison with other measures of confidence

We measured the ability to discriminate correct from incorrect trials (also known as “metacognitive ability”) using an approach inspired by signal detection theory: we computed the type-2 area under the receiver operating characteristic curve (AUROC2)^44^. Given a set of confidence ratings from 1 to 6, there are 5 possible “criterions” to classify trials as having “low” or “high” confidence. The AUROC2 is constructed by measuring the area under the curve defined by the hit rate (i.e., the proportion of correct trials that were reported with high confidence) versus the false alarm rate (i.e., the proportion of incorrect trials that were reported with high confidence).*β*_*p*_ correlated with the AUROC2 (*r*=0.53, *p*<0.01).

We also estimated the participants’ ability to discriminate different conditions. To this end, we obtained the distributions of ratings at each variance condition, and computed their Jensen-Shannon (JS) divergence^11^. The JS divergence between two distributions *P* and *Q* is

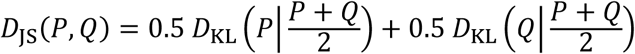
where *D*_KL_(*P*|*Q*) is the Kullback-Leibler^45^ divergence between *P*and *Q*,

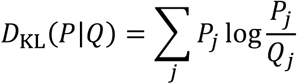

For each participant, we computed the distribution of confidence ratings at each of the four variance levels, and averaged the Jensen-Shannon divergences across all pairs.*β*_*I*_ was correlated with this quantity (*r*=0.86, *p*<10^−8^).

### Statistical analyses

In Experiment 1, we computed the average performance for each variance condition and each participant. These values were submitted to a one-way repeated measures analysis of variance (rm-ANOVA) with factor “variance condition” (4 levels) and “participant” (30 levels) as repeated measure (**Fig. 1**). The normality assumption of this test was checked using the Lilliefors test (*k*=0.7, *c*=0.8, *p*=0.07). We also computed the average confidence rating for each variance condition and each participant, conditioned on correct or incorrect trials, and submitted those values to a two-way rm-ANOVA with factors “variance condition” (4 levels), “outcome” (2 levels: correct or incorrect), and “participant” (30 levels) as repeated measure (**Fig. 2c**). The normality assumption of this test was checked using the Lilliefors test (*k*=0.04, *c*=0.06, *p*>0.5). The goodness of the fit for each model and subject (**Supplementary Fig. 1b**), quantified by the negative log-likelihood (Equation [3]), was submitted to a two-sided paired t-test (29 degrees of freedom). The normality assumption of this test was checked using the Lilliefors test (*k*=0.08, *c*=0.11, *p*>0.5).

In Experiment 2, we compared the within-participants distances in the space defined by (*β*_*p*_,*β*_*I*_ *α*_3_) with the between-subjects distances. Because we have 14 participants, this defines 14 within-subjects distances and 14×13/2=91 between-subjects distances. We z-scored each dimension and used the Euclidean metric to compute distance. The Lilliefors test rejected the null hypothesis that these values were normal (*k*=0.1, *c*=0.08, *p*=0.01); therefore, we used a non-parametric test, the Wilcoxon ranked sum test. This test is unpaired and the reported *p*-value is two-sided.

In Experiment 3, we computed the average performance for each variance condition, task, and participant (**Fig. 5a,b**). We submitted these values to a two-way rm-ANOVA with factors “variance condition” (4 levels), “task” (2 levels), and “participants” (20 levels) as repeated measure. The normality assumption of this test was checked using the Lilliefors test (*k*=0.07, *c*=0.09, *p*=0.36). We computed the average confidence rating across all conditions and participants and performed the same rm-ANOVA used in Experiment 1 (**Fig. 5c,d**). As in Experiment 1, average confidence was normally distributed (Lilliefors test, *k*=0.06, *c*=0.07, *p*=0.17). To evaluate the stability of (*β*_*p*_,*β*_*I*_ *α*_3_) across domains, we computed the within- and between-subjects distances following the same procedure of Experiment 2, and compared these values using the same non-parametric test.

### Data and code availability

The data and codes that support our findings are available upon request to the corresponding author.

---

**Supplementary Figure 1.**
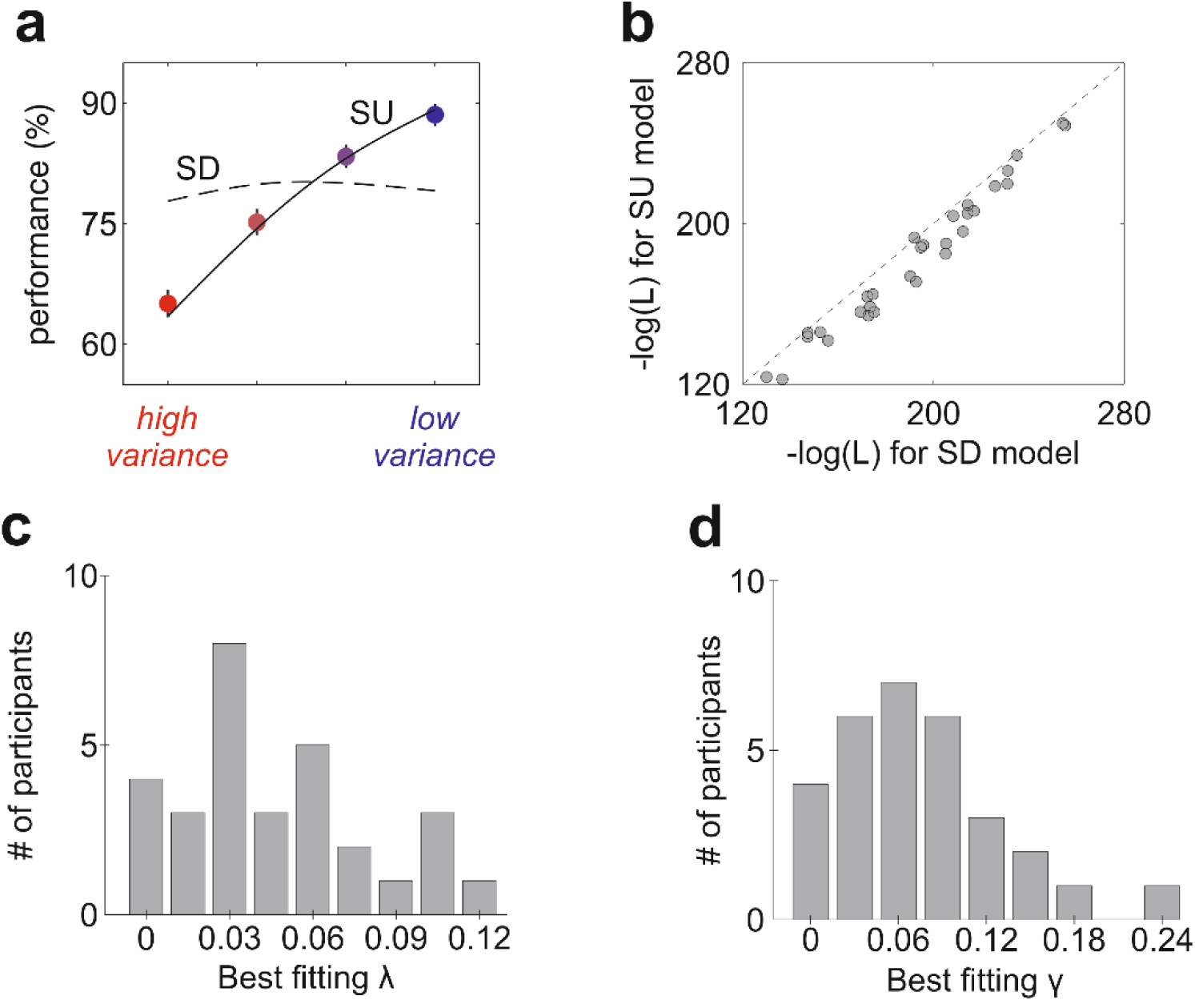
Model fitting results. We fit two probabilistic models that make different assumptions about how decisions are made. The stochastic updating (SU) model is described in the main text (Equations [1] and [2]). In the stochastic decisions (SD) model, the agent makes deterministic updates,

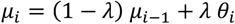

and then makes a softmax decision,

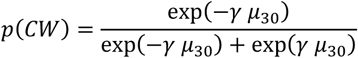

Where *p*(*CW*) is the probability of choosing clockwise and γ is the inverse temperature of the *softmax* rule. In this model, the agent updates perfectly and uses a stochastic (and thus suboptimal) rule for action selection; errors are due to noise in the decisional stage. In the SU model, the updating process is stochastic (Equation [1] in the main text), and decisions are optimal based on the perceived estimate; errors are due to uncertainty in the updating process. Both models fit two parameters (*λ* and *γ*) to the data of each individual. **a**) The SU model (solid line) but not the SD model (dotted line) fits the pattern of increasing performance with decreasing variance. **b)** Model comparison: negative log likelihood of the SU and SD models using the best fitting parameters. Each dot is a different participant. The SU model fits the data significantly better than the SD model (t_(29)_=9.0, p<10^−9^). **c)** Distribution of best fitting parameters *λ* in the SU model across participants. **d)** Distribution of best fitting parameters *γ* in the SU model across participants.

**Supplementary Figure 2.**
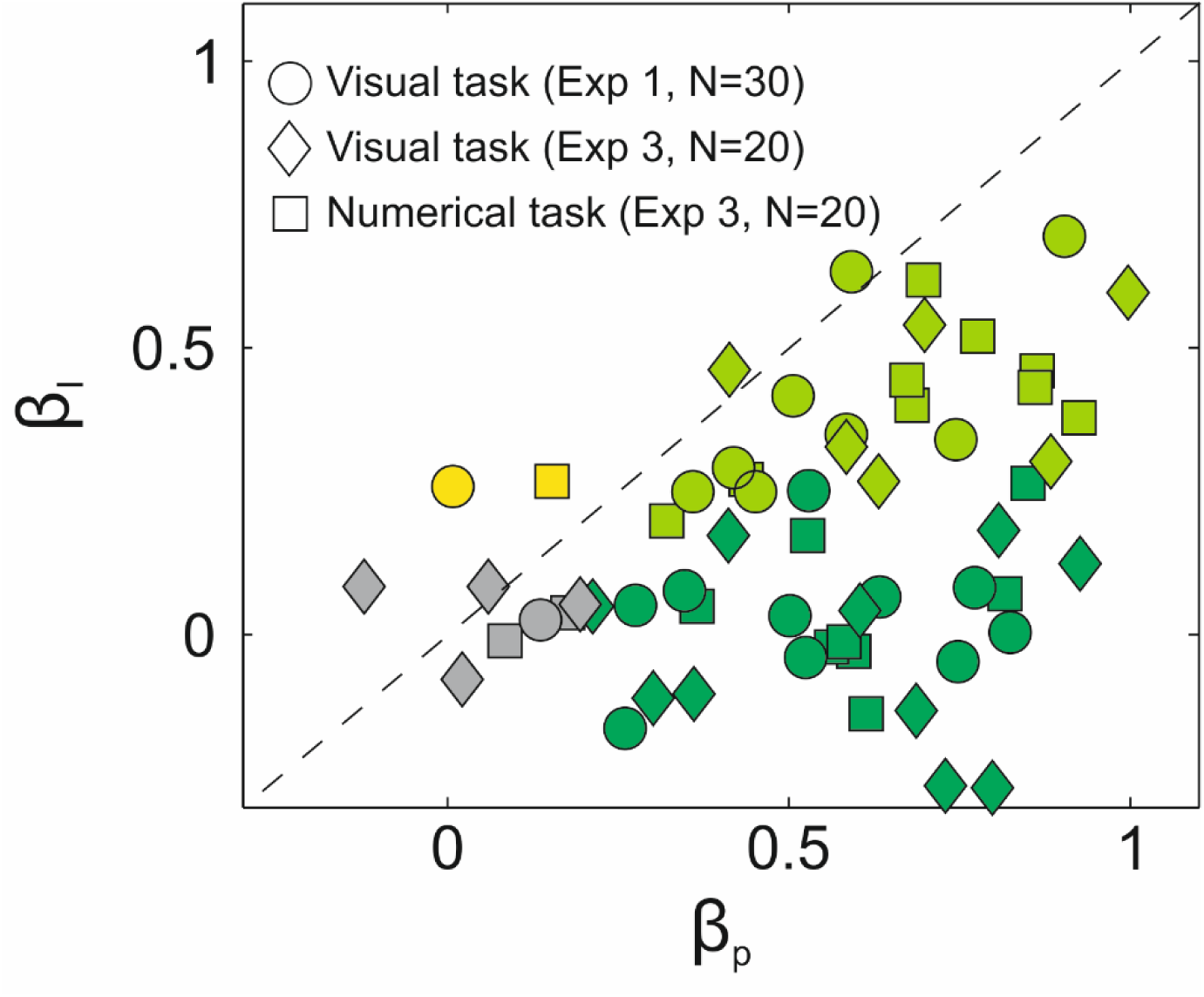
Analysis of confidence across domains. The plot shows regression weights on confidence for different individuals. x-axis: weight of the probability of being correct (*β*_*p*_); y-axis: weight of information (*β*_*I*_). Each marker (circle, diamond, or square) represents one experiment. The colour codes for significance (at the 0.05 level) are as follows: dark green, only *β*_*p*_ was significant; light green, both *β*_*p*_ and *β*_*I*_ were significant; yellow, only *β*_*I*_ was significant; grey, neither was significant. Circles: 30 participants performing the visual task in Experiment 1. Diamonds: 20 other participants performing the visual task in Experiment 3. Squares: the same 20 participants of Experiment 3 performing the numerical averaging task.

**Supplementary Figure 3.**
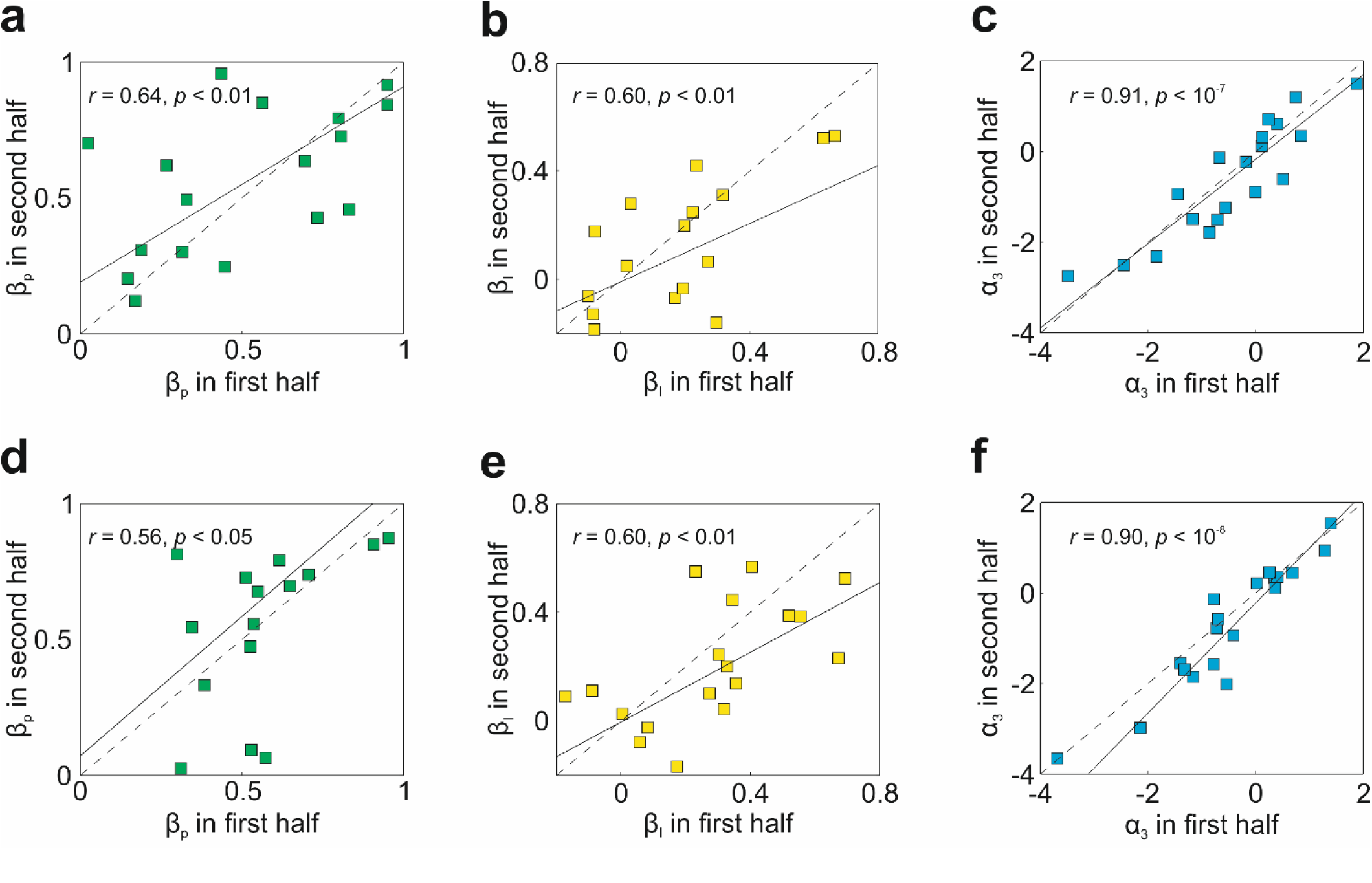
Experiment 3: Stability within each experiment for the visual (**a-c**) and numerical (**d-f**) task. For each half of the experiment (200 trials each), we decomposed confidence in terms of the weight of 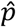(correct) (*β*_*p*_), the weight of information (*β*_*I*_), and the constant term *α*_3_. Correlation across halves for *β*_*p*_ (**a**/**d**), *β*_*I*_ (**b**/**e**), and *α*_3_ (**c**/**f**). Each square is a different participant, the dotted line is the identity, and the value of *r* given in each box is the Pearson correlation coefficient. All three variables are stable within each experiment for both the visual and numerical task.

